# Protonation states in SARS-CoV-2 main protease mapped by neutron crystallography

**DOI:** 10.1101/2020.09.22.308668

**Authors:** Daniel W. Kneller, Gwyndalyn Phillips, Kevin L. Weiss, Swati Pant, Qiu Zhang, Hugh M. O’Neill, Leighton Coates, Andrey Kovalevsky

## Abstract

The main protease (3CL M^pro^) from SARS-CoV-2, the etiological agent of COVID-19, is an essential enzyme for viral replication, possessing an unusual catalytic dyad composed of His41 and Cys145. A long-standing question in the field has been what the protonation states of the ionizable residues in the substrate-binding active site cavity are. Here, we present the room-temperature neutron structure of 3CL M^pro^ from SARS-CoV-2, which allows direct determination of hydrogen atom positions and, hence, protonation states. The catalytic site natively adopts a zwitterionic reactive state where His41 is doubly protonated and positively charged, and Cys145 is in the negatively charged thiolate state. The neutron structure also identified the protonation states of other amino acid residues, mapping electrical charges and intricate hydrogen bonding networks in the SARS-CoV-2 3CL M^pro^ active site cavity and dimer interface. This structure highlights the ability of neutron protein crystallography for experimentally determining protonation states at near-physiological temperature – the critical information for structure-assisted and computational drug design.

## INTRODUCTION

COVID-19, a deadly disease caused by the novel coronavirus SARS-CoV-2 (Severe Acute Respiratory Syndrome Coronavirus 2), is a pandemic of extraordinary proportions, disrupting social life, travel, and the global economy. The development of vaccines and therapeutic intervention measures promises to mitigate the spread of the virus and to alleviate the burdens COVID-19 has caused in many communities in recent months.^1–6^ SARS-CoV-2 is a single-stranded positive-sense RNA virus with a genome comprised of around 30,000 nucleotides. The viral replicase gene encodes two overlapping polyproteins, pp1a and pp1ab, that consist of individual viral proteins essential for replication.^7,8^ Each polyprotein must be processed into individual functional proteins – a vital step in the virus life cycle. This is accomplished by a chymotrypsin-like protease, 3CL M^pro^ or main protease, a hydrolase enzyme that cleaves peptide bonds. The proper functioning of 3CL M^pro^ is indispensable for SARS-CoV-2 replication, whereas its inhibition leads to the inability to produce mature infectious virions. Thus, the enzyme is considered a promising target for the design and development of SARS-CoV-2 specific protease inhibitors and for repurposing existing clinical drugs.^9–15^

SARS-CoV-2 3CL M^pro^ is a cysteine protease^16,17^ and is catalytically active as a homodimer (Figure 1). Its amino acid sequence is 96% homologous to the earlier SARS-CoV 3CL M^pro^, and the catalytic efficiencies of the two enzymes are similar.^10,11,18–20^ The ~34 kDa enzyme has three distinct domains – catalytic domains I (residues 8-101) and II (residues 102-184), and the α-helical domain III (residues 201-303), which is required for protein dimerization.^10,11^ Importantly, the monomeric enzyme shows no catalytic activity, as was demonstrated for SARS-CoV 3CL M^pro^.^21–24^ The catalytic site of 3CL M^pro^ employs a non-canonical Cys145-His41 dyad, thought to be assisted by a water molecule hydrogen bonded to the catalytic histidine.^18,25^ The cysteine thiol group functions as the nucleophile during the first step of the hydrolysis reaction by attacking the carbon atom of the scissile peptide bond. The enzyme recognizes a general amino acid sequence of Leu-Gln↓Ser-Ala-Gly, where ↓ marks the cleavage site, but displays some substrate sequence promiscuity. The active site cavity is located on the surface of the protease and can bind substrate residues in positions P1’ through P5 in the substrate binding subsites S1ʹ through S5ʹ, respectively (Figure 2). Subsites S1, S2 and S4 are shaped into well-formed binding pockets, whereas S1ʹ, S3, and S5 are located on the protein surface with no defined shape. The peptide bond cleavage occurs between the substrate residues at the C-terminal position P1ʹ and N-terminal position P1.

**Figure 1.**
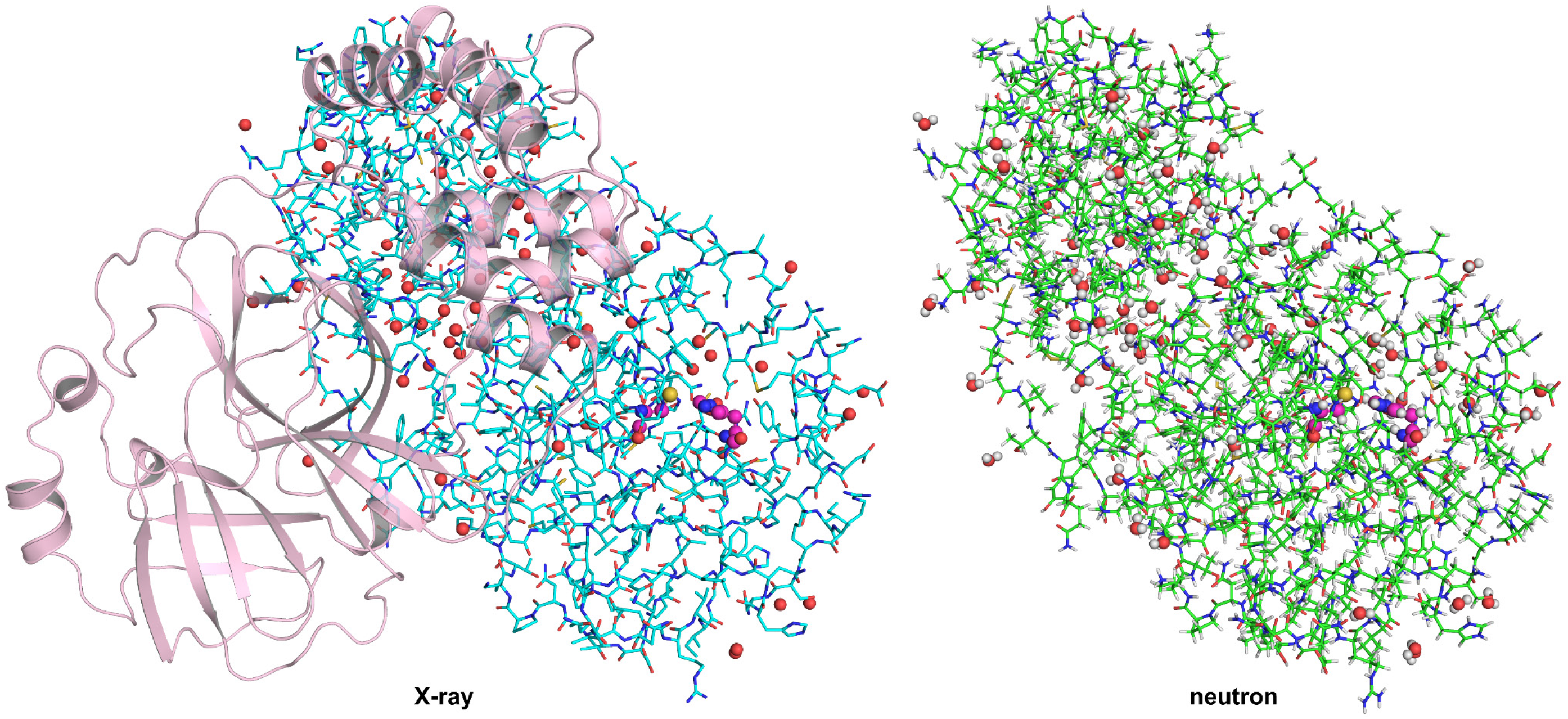
Structure of SARS-CoV-2 3CL M^pro^. The catalytically active dimer is shown on the left. The enzyme monomer shown in stick representation with carbon atoms colored cyan demonstrates that X-ray diffraction only provides the positions of the heavy atoms, i.e. C, N, O, and S. The same monomer shown on the right with carbon atoms colored green illustrates that neutron diffraction provides accurate positions of all atoms, including hydrogen and deuterium. Only oxygen atoms (red spheres) are visible for water molecules in the X-ray structure, whereas all three atoms are fully visible in the neutron structure. The catalytic residues Cys145 and His41 are highlighted in ball-and-stick representation, with carbon atoms shown in magenta.

**Figure 2.**
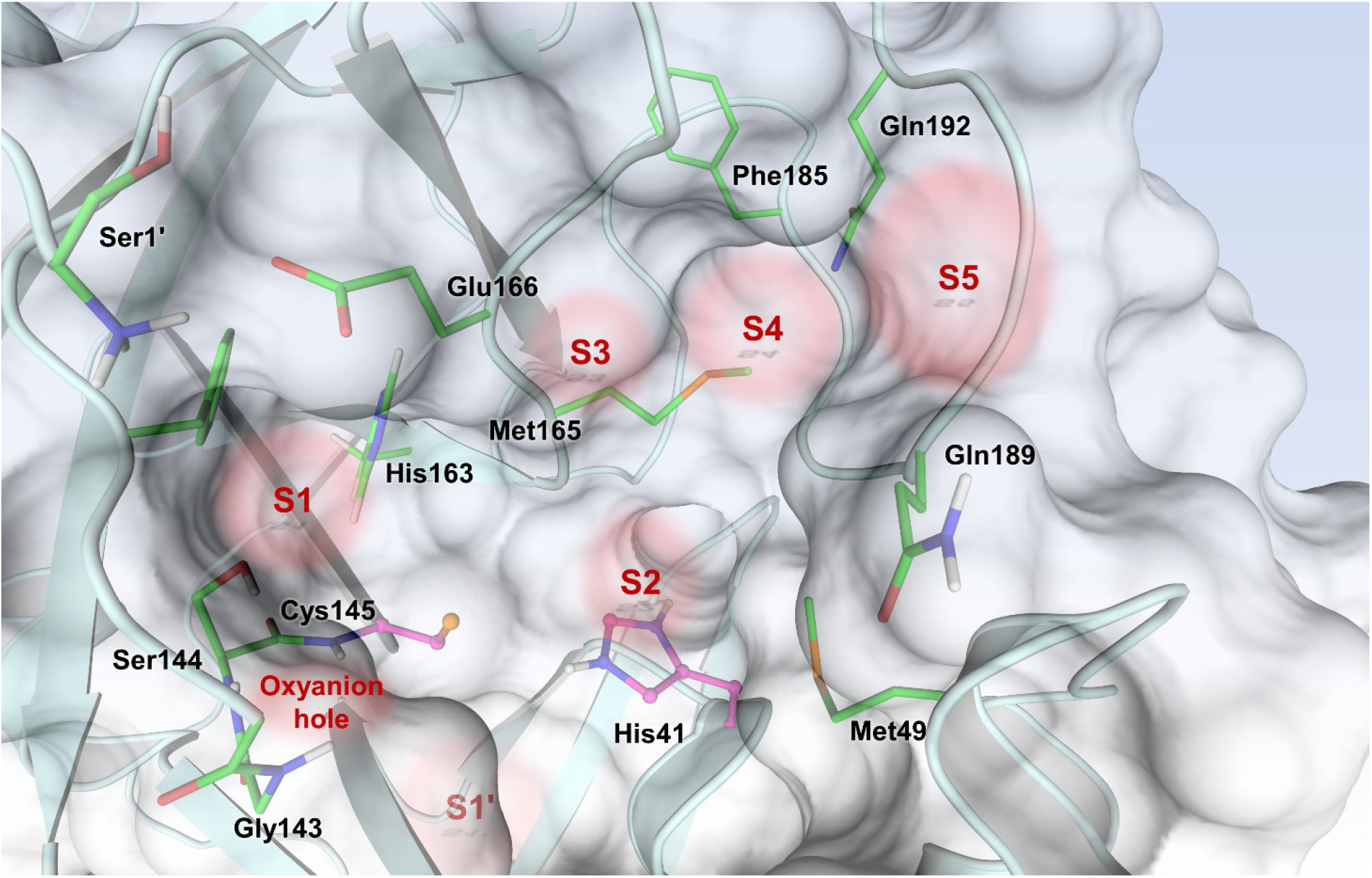
SARS-CoV-2 3CL M^pro^ active site architecture indicating the positions of substrate binding subsites S1’-S5 and the oxyanion hole.

Current structure-assisted drug design efforts are mainly directed towards reversible and irreversible covalent inhibitors that mimic the protease substrate binding to subsites S1ʹ-S5 in the active site cavity,^9–12,14^ whereas the dimer interface can also be explored for the design of dimerization inhibitors.^26,27^ Knowledge of the SARS-CoV-2 3CL M^pro^ active site cavity structure at an atomic level of detail, including the actual locations of hydrogen (H) atoms, can provide critical information to improve rational drug design. The presence or absence of H atoms at specific sites on amino acid residues determines their protonation states and, thus, their electrical charges, defining the electrostatics and hydrogen bonding interactions. Of note, half of all atoms in protein and small-molecule drugs are H. X-ray crystallography is typically the standard experimental method for structure-assisted drug design but is unable to reliably locate H atoms in biological macromolecules, leaving significant gaps in our understanding of biological function and drug binding.^28^ Electron clouds scatter X-rays, thus scattering power is determined by the number of electrons present in an atom, *i.e.* by its atomic number. H, with just a single electron that often participates in highly polarized chemical bonds, is the weakest possible X-ray scatterer and consequently is invisible in X-ray structures with a few exceptions beyond subatomic resolution.^29–31^ In contrast, atomic nuclei scatter neutrons, where the scattering power of neutrons is independent of the atomic number. Deuterium (D), a heavy isotope of hydrogen (H), scatters neutrons just as well as carbon, nitrogen, and oxygen. Neutron crystallography is capable of accurately determining positions of H and D atoms and visualizing hydrogen bonding interactions at moderate resolutions,^32–35^ where X-rays cannot locate the functional H atoms.^29^ Moreover, unlike X-rays,^36^ neutrons cause no direct or indirect radiation damage to protein crystals, permitting diffraction data collection at near-physiological (room) temperature and avoiding possible artifacts induced by the use of cryoprotectant chemicals required for X-ray cryo-crystallographic measurements.

We produced neutron-quality crystals of the ligand-free SARS-CoV-2 3CL M^pro^ at pH 6.6 and obtained a room-temperature neutron structure of the enzyme refined jointly with a room-temperature X-ray dataset collected from the same crystal (Figure 1).^37^ We accurately determined the locations of exchangeable H atoms that were observed as Ds attached to electronegative atoms such as oxygen, nitrogen, or sulfur atoms. Our experimental observations identified the protonation states of ionizable amino acid residues allowing us to accurately map hydrogen bonding networks in the SARS-CoV-2 3CL M^pro^ active site cavity and throughout the enzyme structure. Neutron diffraction data were collected from an H/D-exchanged SARS-CoV-2 3CL M^pro^ crystal at pD 7.0 (pD = pH +0.4) to 2.5 Å resolution. In neutron crystallographic experiments, protein crystals are usually H/D exchanged with the heavy water (D_2_O) to increase the signal-to-noise ratio of the diffraction pattern as H has a large incoherent scattering cross section that increases background. Also, the coherent neutron scattering length of H is negative (−3.739 fm, (https://www.ncnr.nist.gov/resources/n-lengths/) and are therefore observed in the neutron scattering length (or nuclear) density maps as troughs. At moderate resolutions, the negative neutron scattering length of H leads to the density cancellation phenomenon observed for CH, CH_2_ and CH_3_ groups as H atoms attached to carbon atoms cannot exchange with D. Conversely, D has a coherent neutron scattering length of +6.671 fm and, thus, is observed as peaks in nuclear density maps. Because D atoms scatter neutrons just as well as other protein atoms, they can be directly detected in neutron structures at moderate resolutions as low as 2.5-2.6 Å.^38,39^ Notably, sulfur (S) has a coherent neutron scattering length of +2.847 fm, less than half the magnitude of that for carbon, oxygen, nitrogen, and deuterium. Consequently, deprotonated thiol groups (S^−^) in cysteine (Cys) and side chain S atoms in methionine (Met) residues are often not visible in nuclear density maps.

## RESULTS

### Protonation states of the catalytic site and nearby residues

The electron density for the catalytic site and the nearby residues of SARS-CoV-2 3CL M^pro^ is shown in Figure 3A. Although hydrogen bonding interactions can be inferred from the distances between the heavy atoms, the locations of H atoms, and the protonation states of the amino acid residues can only be assumed. Instead, the nuclear density map shown in Figure 3B displays the actual positions of exchanged D atoms, accurately visualizing hydrogen bond donors and acceptors. In the neutron structure, we observed that the catalytic Cys145 thiol is in the deprotonated negatively charged thiolate state.

**Figure 3.**
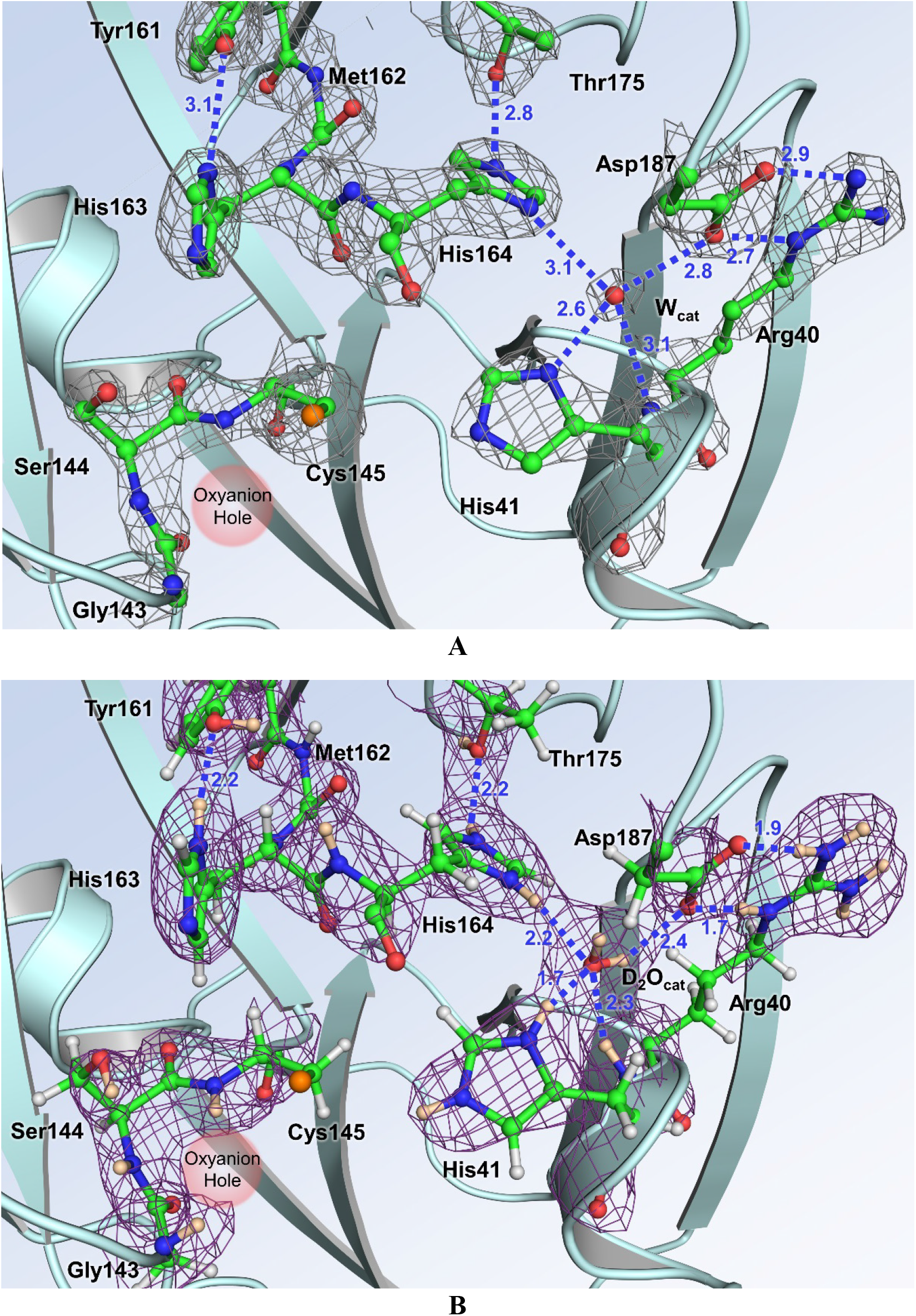
The catalytic site of SARS-CoV-2 3CL M^pro^. (**A**) The 2FO-FC electron density map contoured at 2.0 σ level (grey mesh) with no hydrogen atoms visible. Distances between the heavy atoms in Ångstroms illustrate possible hydrogen bonds. (**B**) The 2FO-FC nuclear density map contoured at 2.0 σ level (violet mesh), allowing visualization of the actual protonation states and hydrogen bonding interactions (D…O distances are shown in Ångstroms).

In contrast, located 3.9 Å away from Cys145, the catalytic residue His41 is protonated on both Nδ1 and Nε2 nitrogen atoms of the imidazole side chain and is therefore positively charged (Figure S1). As a result, the catalytic site natively adopts the zwitterionic reactive state required for catalysis.^19,20,41^ His41 is strongly hydrogen bonded to a water molecule (D_2_O_cat_) that presumably plays the role of the third catalytic residue^18,25^ from a canonical catalytic triad, stabilizing the charge and position of the His41 imidazolium ring. The Nδ1-D…O_D2O_ distance is 1.7 Å. The position of the D_2_O_cat_ molecule is stabilized by several more, but possibly weaker, hydrogen bonds with His41 main chain, His164, and Asp187. His164 is doubly protonated and positively charged; it donates a D in a hydrogen bond with the Thr175 hydroxyl within the interior of the protein. As expected, Asp187 is not protonated and is negatively charged, and participates in a strong salt bridge with Arg40. His163 positioned near the catalytic Cys145 is singly protonated and uncharged, making a hydrogen bond with the phenolic side chain of Tyr161 (Figure S1). The main chain D atoms of Gly143, Ser144, and Cys145 comprising the oxyanion hole are also readily visible in the nuclear density (Figure 3B).

### Substrate binding subsite S1

The P1 group of a substrate, usually Gln, binds in a rather peculiar substrate binding subsite S1 (Figure 4A). From one side, it is flanked by residues 140-144, making a turn that creates the oxyanion hole and, on the opposite side, by Met165, Glu166, and His172. The back wall of subsite S1 is created by the side chains of Phe140 and His163. Interestingly, the N-terminus of Ser1ʹ of the second monomer within the active 3CL M^pro^ homodimer reaches in to cap the subsite S1 from the top. In our neutron structure, the N-terminal amine is the protonated, positively charged −ND_3_+ ammonium cation. It forms three hydrogen bonds, one each with the main chain carbonyl of Phe140, the side chain carboxylate of Glu166, and a D_2_O molecule. Both histidine residues, His163 and His172, in this subsite are singly protonated and neutral (Figure S1). The deprotonated Nδ1 of His172 is hydrogen bonded with the main chain amide nitrogen of Gly138 with an N…D distance of 2.2 Å, whereas the protonated Nε2 makes a long, possibly weak, hydrogen bond interaction with the Glu166 carboxylate with the D…O distance of 2.5 Å.

**Figure 4.**
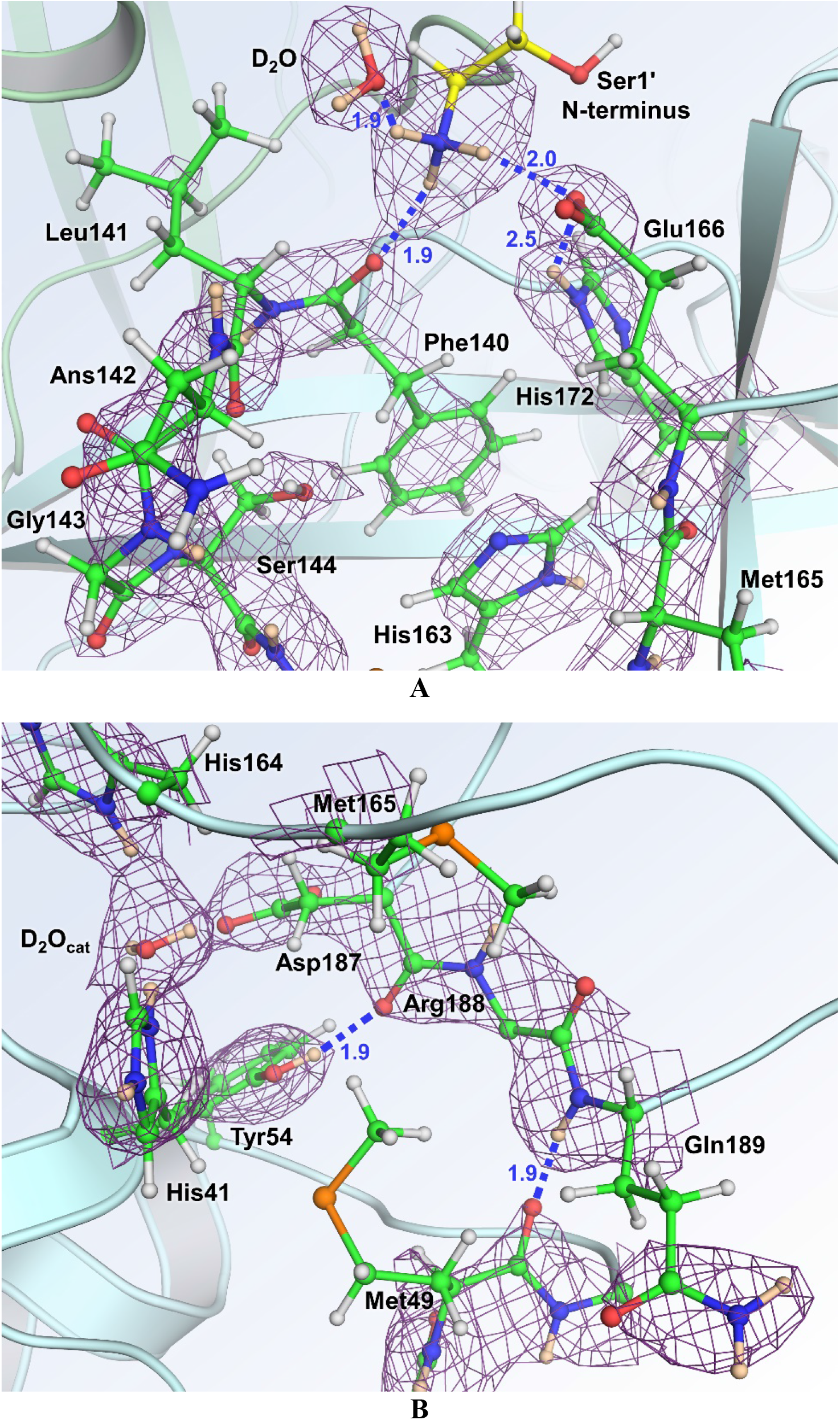
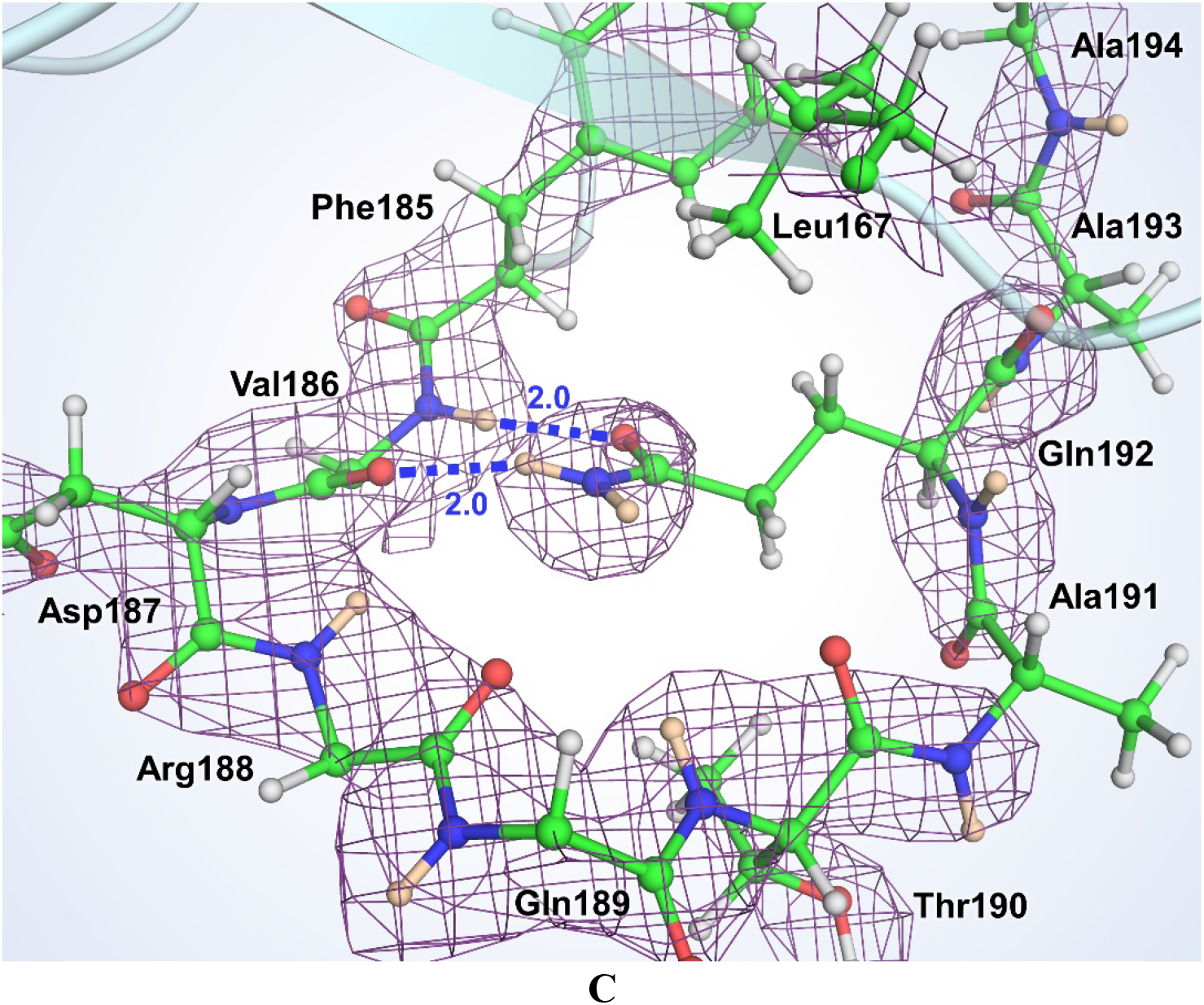
The 2FO-FC nuclear density map contoured at 2.0 σ level (violet mesh) for the residues in the substrate binding subsites S1 (**A**), S2 (**B**) and S4 (**C**). Hydrogen bonds are shown as blue dashed lines. The D…O distances are in Ångstroms. For clarity, the side chains were omitted for Val186, Arg188 and Gln189.

### Substrate binding subsite S2

Subsite S2 is more hydrophobic than subsite S1, as it binds the hydrophobic residues Leu or Phe in substrate position P2 (Figure 4B). Subsite S2 is flanked by the π-systems of His41 and the main chains connecting Asp187, Arg188, and Gln189. It is capped by Met165 whereas Met49, situated on the short P2 helix spanning residues Ser46 through Leu50, adopts a conformation impeding the entrance to this site. Met49 appears to be conformationally flexible, vacating its position in the ligand-free enzyme to allow various P2 groups to occupy this subsite when inhibitors bind.^9–11^ Tyr54 serves as the base of subsite S2, and its phenolic hydroxyl group donates its D in a hydrogen bond with the main chain carbonyl of Asp187.

### Substrate binding subsites S3-S5

Among the substrate binding subsites S3, S4, and S5, only subsite S4 has a well-defined binding pocket architecture. Subsites S3 and S5 are on the protein surface, fully exposed to the bulk solvent, and have ill-defined borders. Subsite S3 is located between residues Glu166 and Gln189 that are >9 Å apart. Subsite S5 is between Pro168 of the β-hairpin flap, spanning residues Met165 through His172, and the P5 loop consisting of residues Thr190 through Ala194 (Figure 2). Subsite S4 is formed between a long loop spanning residues Phe185 through Ala194 that acts as its base and the β-hairpin flap on the top. The loop turns 180° at Gln189, with its secondary structure being stabilized by hydrogen bonds between the Gln192 side chain amide and the main chain atoms of Val186. The side chains of Leu167 and Phe185 form the back wall of this site in the protein interior, creating a deep, mainly hydrophobic pocket (Figure 4C).

### Dimer interface

The two protomers in the SARS-CoV-2 3CL M^pro^ homodimer interact through an extensive dimer interface. Protomer 1 (unprimed residue numbers) forms elaborate networks of hydrogen bonding interactions with N-terminal residues 1ʹ-16ʹ, a β-strand with residues 118ʹ-125ʹ, and a loop containing residues 137ʹ-142ʹ of protomer 2 (primed residue numbers, Figure 5). There are also many hydrophobic interactions within the dimer interface. The N-termini of the two protomers meet at the start of a short α-helix spanning residues Gly11-Cys16 and Gly11ʹ-Cys16ʹ to form several hydrogen bonds involving the main chain and side chain atoms of Ser10, Gly11, Ser10ʹ, Gly11ʹ and Glu14ʹ (Figure S2A). The N-terminal loop then extends across the face of the opposite protomer to subsite S1. Here, Ala7ʹ, Phe8ʹ, Arg4ʹ, and Ser1ʹ are observed making hydrogen bonds with Val125, Glu290, Lys137, Phe140, and Glu166. Ser123ʹ and Asn119ʹ of protomer 2 β-strand containing residues 118ʹ-125ʹ have direct hydrogen bonds and water-mediated interactions with the C-terminal residues of protomer 1, in addition to the reciprocal hydrogen bonds between Val125’ and Ala7 formed due to the two-fold symmetry of the enzyme dimer (Figure S2B). Similar reciprocity of hydrogen bonding is found for the loop consisting of residues 137ʹ-144ʹ in protomer 2 that interacts with the N-terminal residues Ser1 and Arg4 of monomer 1. Also, the side chain hydroxyl of Ser139ʹ makes a hydrogen bond with the side chain amide of Gln299 (Figure S2C).

**Figure 5.**
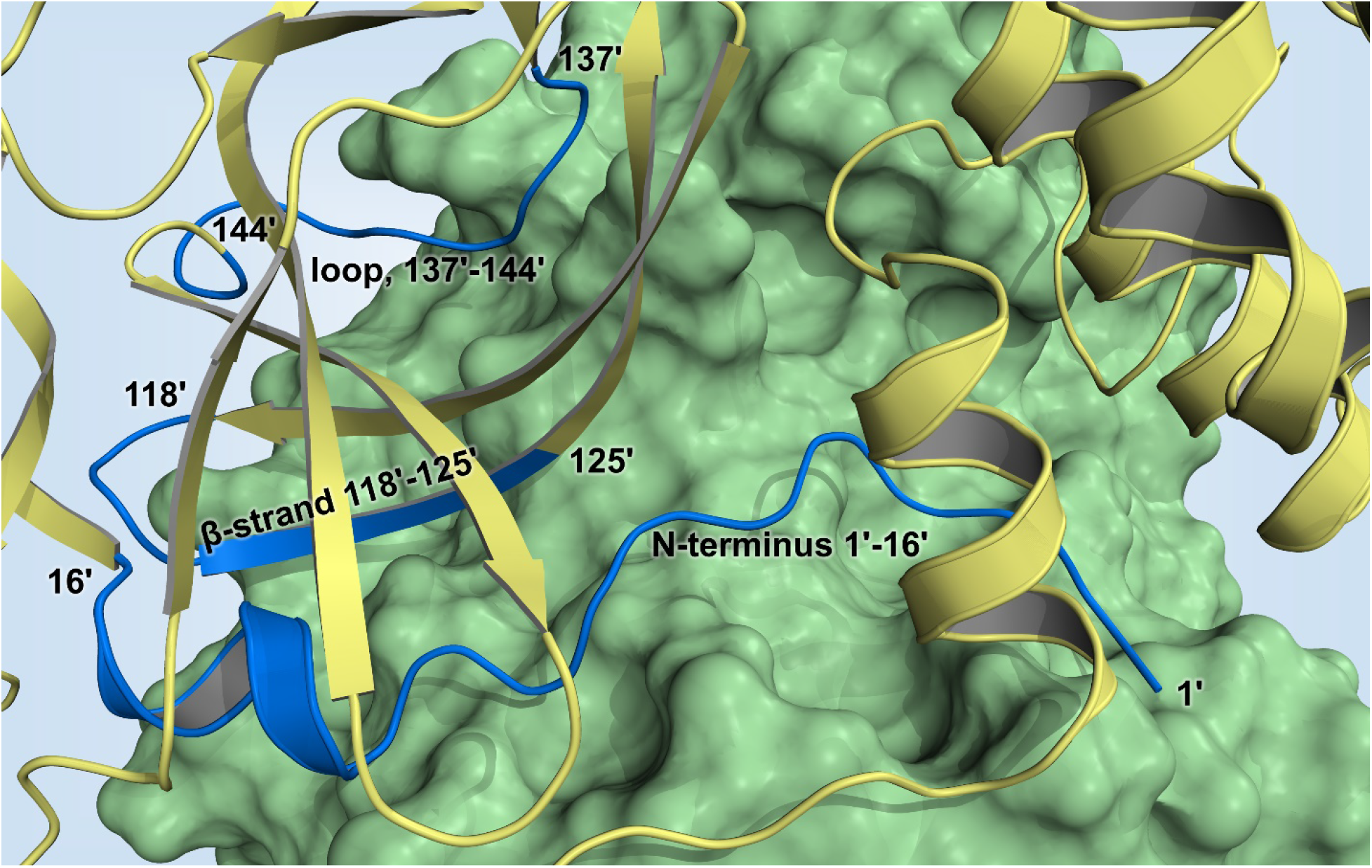
The SARS-CoV-2 3CL M^pro^ dimer interface. Protomer 1 (unprimed residues 1-306) is shown as a green surface, protomer 2 (primed residues 1’-306’) is in cartoon representation colored yellow. The parts of protomer 2 interacting with protomer 1 are colored blue.

### Protonation states of Cys residues

The SARS-CoV-2 3CL M^pro^ structure contains twelve Cys residues spread throughout the sequence. The reason for such a large number of cysteines within the enzyme sequence and their functional roles are unknown, except for the catalytic Cys145. By examining the nuclear density, we were able to establish the protonation states of all Cys residues in the structure (Figure S3 and S4). The side chains of residues Cys16, Cys85, Cys117, Cys156, Cys160, Cys265, and Cys300, are protonated thiol groups. Conversely, the remaining Cys residues – Cys22, Cys38, Cys44, Cys128 and Cys145 – contain deprotonated thiolates. Accordingly, because we can unambiguously differentiate between seven protonated and five deprotonated Cys side chains, this provides strong evidence that the catalytic Cys145 is in the reactive deprotonated negatively charged thiolate form.

## DISCUSSION

Neutron crystallography is the only technique capable of accurately determining the positions of H (and D) atoms in biological macromolecules without causing radiation damage to the samples at near-physiological temperature. We have succeeded in determining a room-temperature neutron structure of the homodimeric SARS-CoV-2 3CL M^pro^ enzyme mapping the protonation states of amino acid residues in the active site cavity and throughout the protein structure. The neutron structure also reveals the elaborate hydrogen bonding networks formed in the catalytic site, including those in the six substrate binding subsites, and throughout the dimer interface. Protonation states are determined by the locations of H atoms, observed in the neutron structure as Ds after the H/D exchange was performed. Also, protonation states establish the electrical charges and therefore determine the electrostatic environment in the protein, essential information for the structure-assisted and computational drug design. The catalytic dyad is observed in the reactive zwitterionic state in the crystal at pD 7.0, having a deprotonated negatively charged Cys145 and doubly protonated positively charged His41, instead of the catalytically resting (non-reactive) state^40^ in which both catalytic residues are neutral (*i.e.*, protonated Cys145 and singly protonated His41). His41 is hydrogen bonded to the catalytic water molecule that is held in place by interactions with the doubly protonated His164 and negatively charged Asp87. The substrate binding subsite S1 contains the protonated positively charged N-terminal amine from protomer 2 and neutral His163. The non-protonated Nε2 of His163 is exposed and can be utilized as a hydrogen bond acceptor by an inhibitor P1 substituent. In the hydrophobic subsite S2, the Tyr54 hydroxyl can act as a hydrogen bond acceptor for a P2 substituent with a hydrogen bond donor functionality. Consequently, specific protease inhibitor design needs to consider the observed charge distribution in the SARS-CoV-2 3CL M^pro^ active site cavity.

We also accurately mapped the hydrogen bonding networks within the dimer interface of SARS-CoV-2 3CL M^pro^. Significantly, the N-terminal residues between Ser1 and Cys16 (and Ser1’ and Cys16’) account for the most hydrogen bonds between the two protomers. There are thirteen direct hydrogen bonds formed by each N-terminus interacting with the opposite protomer. Ser1 is an essential residue for building the correct architecture of substrate binding subsite S1. Truncation of the first four N-terminal residues in SARS-CoV 3CL M^pro^ has previously shown to disrupt dimerization and dramatically diminish enzyme activity.^22^ We, therefore, suggest that the enzyme area that interacts with residues 1 through 16 from the other protomer may be the most promising part of the dimer interface for designing SARS-CoV-2 3CL M^pro^-specific dimerization inhibitors, as was also proposed for SARS-CoV enzyme.^41,42^

## CONCLUSION

We have successfully determined the neutron structure of the ligand-free SARS-CoV-2 3CL M^pro^ at near-physiological (room) temperature and mapped the locations of H atoms (observed as D) in the enzyme. Thus, protonation states and electrical charges of the ionizable residues have been accurately resolved. Establishing the electrostatics in the enzyme active site cavity and at the dimer interface is critical information to guide the structure-assisted and computational drug design of protease inhibitors, specifically targeting the enzyme from SARS-CoV-2.

## METHODS

### General Information

Protein purification columns were purchased from Cytiva (Piscataway, New Jersey, USA). Crystallization reagents were purchased from Hampton Research (Aliso Viejo, California, USA). Crystallographic supplies were purchased from MiTeGen (Ithaca, New York, USA) and Vitrocom (Mountain Lakes, New Jersey, USA).

### Cloning, expression, and purification of SARS-CoV-2 3CL M^pro^

The 3CL M^pro^ (Nsp5 M^pro^) from SARS-CoV-2 was cloned into pD451-SR plasmid harboring kanamycin resistance (ATUM, Newark, CA), expressed and purified according to the procedure published elsewhere.^25^ To make the authentic the N-terminus of 3CL M^pro^, the enzyme sequence is flanked by the maltose binding protein (MBP) followed by the protease autoprocessing site SAVLQ↓SGFRK (down arrow indicates the autocleavage site) which corresponds to the cleavage position between NSP4 and NSP5 in the viral polyprotein. To form the authentic C-terminus, the enzyme construct codes for the human rhinovirus 3C (HRV-3C) protease cleavage site (SGVTFQ↓GP) connected to a 6xHis tag. The N-terminal flanking sequence is autocleaved during the expression in *E. coli* (BL21 DE3), whereas the C-terminal flanking sequence is removed by the treatment with HRV-3C protease (Millipore Sigma, St. Louis, MO). For crystallization, the authentic 3CL M^pro^ is concentrated to ~4 mg/mL in 20 mM Tris, 150 mM NaCl, 1 mM TCEP, pH 8.0.

### Crystallization

The methodology for growing large crystals of 3CL M^pro^ suitable for neutron diffraction is described as follows. Initial protein crystallization conditions were discovered by screening conducted at the Hauptman-Woodward Medical Research Institute (HWI).^43^ Crystal aggregates were reproduced using the sitting drop vapor diffusion method using 25% PEG3350, 0.1 M Bis-Tris pH 6.5. Microseeding using Hampton Research Seed Beads™ was performed to grow neutron-quality crystals in Hampton 9-well plates and sandwich box set-ups with 200 μL drops of protein mixed with 18% PEG3350, 0.1 M Bis-Tris pH 6.0, 3% DMSO at a 1:1 ratio seeded with 1 μL of microseeds at a 1:500 dilution. This condition produced a final pH in the crystallization drop of 6.6 as measured by microelectrode. The crystal tray used to harvest a crystal for neutron diffraction was incubated initially at 18°C then gradually lowered to 10°C over several weeks. The crystal grew to final dimensions of ~2×0.8×0.2 mm (~0.3 mm^3^) in a triangular plate-like morphology. The crystal was mounted in a quartz capillary accompanied with 18% PEG3350 prepared with 100% D_2_O and allowed to H/D exchange for several days before starting the neutron data collection.

### Neutron diffraction data collection

The large crystal was screened for diffraction quality using a broad-bandpass Laue configuration using neutrons from 2.8 to 10 Å at the IMAGINE instrument at the High Flux Isotope Reactor (HFIR) at Oak Ridge National Laboratory.^44–46^ Neutron diffraction data were collected using the Macromolecular Neutron Diffractometer (MaNDi) instrument at the Spallation Neutron Source (SNS).^45,47-49^ The crystal was held still at room temperature, and diffraction data were collected for 24 hours using all neutrons between 2-4 Å. Following this, the crystal was rotated by Δϕ = 10°, and a subsequent data frame was collected again for 24 hours. A total of twenty-three data frames were collected in the final neutron dataset. Diffraction data were reduced using the Mantid package, with integration carried out using three-dimensional TOF profile fitting.^50^ Wavelength normalization of the Laue data was performed using the Lauenorm program from the Lauegen suite.^51,52^ The neutron data collection statics are shown in Table S1.

### X-ray diffraction data collection

The room-temperature diffraction dataset was collected from the same crystal used for the neutron data collection using a Rigaku HighFlux HomeLab instrument equipped with a MicroMax-007 HF X-ray generator and Osmic VariMax optics. The diffraction images were obtained using an Eiger R 4M hybrid photon counting detector. Diffraction data were integrated using the CrysAlis Pro software suite (Rigaku Inc., The Woodlands, TX). Diffraction data were then reduced and scaled using the Aimless^53^ program from the CCP4 suite^54^; molecular replacement using PDB code 6WQF^25^ was then performed with Molrep^54^ from the CCP4 program suite. The protein structure was first refined against the X-ray using *Phenix.refine* from the Phenix^55^ suite of programs and the *Coot*^56^ molecular graphics program to obtain an accurate model for the subsequent X-ray/neutron joint refinement. The geometry of the final model was then carefully checked with Molprobity.^57^ The X-ray data collection statics are shown in Table S1.

### Joint X-ray/neutron refinement

The joint X-ray/neutron refinement of ligand-free 3CL M^pro^ was performed using *nCNS*^58,59^ and the structure was manipulated in *Coot*.^56^ After initial rigid-body refinement, several cycles of positional, atomic displacement parameter, and occupancy refinement were performed. The structure was checked for the correctness of side-chain conformations, hydrogen bonding, and orientations of D_2_O water molecules, which were built based on the mFo-DFc difference neutron scattering length density maps. The 2mFo-DFc and mFo-DFc neutron scattering length density maps were then examined to determine the correct orientations of hydroxyl (Ser, Thr, Tyr), thiol (Cys) and ammonium (Lys) groups, and protonation states of the enzyme residues. The protonation states of some disordered side chains could not be obtained directly and remained ambiguous. All water molecules were refined as D_2_O. Initially, water oxygen atoms were positioned according to their electron density peaks and then were shifted slightly in accordance with the neutron scattering length density maps. All labile H positions in 3CL M^pro^ were modeled as D atoms, and then the occupancies of those atoms were refined individually within the range of −0.56 (pure H) to 1.00 (pure D). Before depositing the neutron structure to the PDB, a script was run that converts a record for the coordinates of a D atom into two records corresponding to an H and a D partially occupying the same site, both with positive partial occupancies that add up to unity. The percent D at a specific site is calculated according to the following formula: %D = {Occup.(D) + 0.56}/1.56.

## Data availability

The coordinates and structure factors for ligand-free SARS-CoV-2 3CL Mpro have been deposited in the PDB with accession code 7JUN. Any other relevant data are available from the corresponding authors upon reasonable request.

## Acknowledgements

This research was supported by the DOE Office of Science through the National Virtual Biotechnology Laboratory (NVBL), a consortium of DOE national laboratories focused on response to COVID-19, with funding provided by the Coronavirus CARES Act. This research used resources at the Spallation Neutron Source and the High Flux Isotope Reactor, which are DOE Office of Science User Facilities operated by the Oak Ridge National Laboratory. The Office of Biological and Environmental Research supported research at ORNL’s Center for Structural Molecular Biology (CSMB), a DOE Office of Science User Facility. This research used resources at the Second Target Station, which is a DOE Office of Science User Facilities Construction Project at Oak Ridge National Laboratory.

## Author contributions

L.C and A.K. conceived the study. A.K., G.P. and Q.Z. designed and cloned the gene. K.L.W. and H.M.O’N. contributed to experimental design and implementation of protein expression. G.P., S.P., and Q.Z. expressed and purified the protein, D.W.K. and A.K. crystallized the protein. A.K. and D.W.K. collected the X-ray diffraction data. L.C. collected and reduced the neutron diffraction data. A.K., D.W.K. and L.C. refined the structure. D.W.K., L.C., and A.K. wrote the paper with help from all co-authors.

